# The landscape of viral associations in human cancers

**DOI:** 10.1101/465757

**Authors:** Marc Zapatka, Ivan Borozan, Daniel S. Brewer, Murat Iskar, Adam Grundhoff, Malik Alawi, Nikita Desai, Holger Sültmann, Holger Moch, PCAWG Pathogens Working Group, ICGC/TCGA Pan-cancer Analysis of Whole Genomes Network, Colin S. Cooper, Roland Eils, Vincent Ferretti, Peter Lichter

## Abstract

Potential viral pathogens were systematically investigated in the whole-genome and transcriptome sequencing of 2,656 donors as part of the Pan-Cancer Analysis of Whole Genomes using a consensus approach integrating three independent pathogen detection pipelines. Viruses were detected in 382 genomic and 68 transcriptome data sets. We extensively searched and characterized numerous features of virus-positive cancers integrating various PCAWG datasets. We show the high prevalence of known tumor associated viruses such as EBV, HBV and several HPV types. Our systematic analysis revealed that HPV presence was significantly exclusive with well-known driver mutations in head/neck cancer. A strong association was observed between HPV infection and the APOBEC mutational signatures, suggesting the role of impaired mechanism of antiviral cellular defense as a driving force in the development of cervical, bladder and head neck carcinoma. Viral integration into the host genome was observed for HBV, HPV16, HPV18 and AAV2 and associated with a local increase in copy number variations. The recurrent viral integrations at the *TERT* promoter were coupled to high telomerase expression uncovering a further mechanism to activate this tumor driving process. High levels of endogenous retrovirus ERV1 expression is linked to worse survival outcome in kidney cancer.

---

The World Health Organization estimates that 15.4% of all cancers are attributable to infections and 9.9% are linked to viruses^1, 2^. Cancers attributable to infections have a greater incidence than any individual type of cancer worldwide. Eleven pathogens have been classified as carcinogenic agents in humans by the International Agency for Research on Cancer (IARC)^3^. After Helicobacter pylori (associated with 770,000 cases), the four most prominent infection related causes of cancer are estimated to be viral^2^: human papilloma virus (HPV)^4, 5^ (associated with 640,000 cancers), hepatitis B virus (HBV)^5^ (420,000), hepatitis C virus (HCV)^6^ (170,000) and Epstein-Barr Virus (EBV)^7^ (120,000). It has been shown that viruses can contribute to the biology of multistep oncogenesis and are implicated in many of the hallmarks of cancer^8^. Most importantly, the discovery of links between infection and cancer types has provided actionable opportunities, such as HPV vaccines as preventive measure, to reduce the global impact of cancer. The following characteristics were proposed to define human viruses causing cancer through direct or indirect carcinogenesis^9^: i) Presence and persistence of viral DNA in tumor biopsies; ii) Growth promoting activity of viral genes in model systems; iii) Dependence of malignant phenotype on continuous viral oncogene expression or modification of host genes; iv) Epidemiological evidence that a virus infection represents a major risk for development of cancer.

The worldwide efforts of comprehensive genome and transcriptome analyses of tissue samples from cancer patients generate congenial facilities for capturing information not only from human cells, but also from other - potentially pathogenic - organisms or viruses present in the tissue. By far the most comprehensive collection of whole genome and transcriptome data from cancer tissues has been generated within the ICGC (International Cancer Genome Consortium) project PCAWG (Pan-Cancer Analysis of Whole Genomes)^10^ providing a unique opportunity for a systematic search for tumor-associated viruses.

The PCAWG working group “Exploratory Pathogens” searched the whole genome sequencing (WGS) and whole transcriptome sequencing (RNA-seq) data of the PCAWG consensus cohort. Focusing on viral pathogens, we applied three independently developed pathogen detection pipelines ‘Computational Pathogen Sequence Identification’ (CaPSID)^11^, ‘Pathogen Discovery Pipeline’ (P-DiP), and ‘SEarching for PATHogens’ (SEPATH) to generate a large compendium of viral associations across 39 cancer types. We extensively characterized the known and novel viral associations by integrating driver mutations, mutational signatures, gene expression profiles and patient survival data of the same set of tumors analyzed in PCAWG.

## Results & Discussion

### Identification of tumor-associated viruses using whole genome and transcriptome sequencing data

To identify the presence of viral sequences, we explored the WGS data of 5,354 tumor/normal samples across 39 cancer types, and 1,057 tumor RNA-seq data across 25 cancer types (Supplementary Table 1, sheet “Candidate Reads WGS” and “Candidate Reads RNAseq”). 195.8 billion reads were considered for further analysis as they were not sufficiently aligned to the human reference genome in the PCAWG-generated alignment (see Materials and Methods). Remaining reads ranged from 28,036 to 800 million per WGS tumor sample and up to 120 million per RNA-seq tumor sample (Figure 1a, Supplementary Figure 1a, b). Viral sequences were detected and quantified independently by three recently developed pathogen discovery pipelines CaPSID, P-DiP and SEPATH (see Supplementary Methods). The estimated relative abundance of a virus was calculated as viral reads per million extracted reads (PMER) at the genus level to improve data consistency between pipelines. To minimize the rate of false positive scores in virus detection, we applied a strict threshold of PMER>1 supported by at least three viral reads as similarly suggested by previous studies^11, 12^. If a viral genus was identified by at least two of the three pipelines, it was considered present as a consensus hit in the sample. In total, 532 genera were considered for the extensive virus search in at least two of the pipelines (Supplementary Figure 1C). Filtering of suspected viral laboratory contaminants was achieved through P-DiP, by examining each assembled contig of viral sequence segments for artificial, non-viral vector sequences and inspecting virus genome coverage across all positive samples (see Materials and Methods). The most frequent hits prone to suspected contamination were lambdavirus, alphabaculovirus, microvirus, simplexvirus, hepacivirus, cytomegalovirus, orthopoxvirus and punalikevirus these were observed across many tumor types (Figure 1b). As a second measure to identify spurious virus detections, we analysed the genome coverage across all virus positive samples (Supplementary Figure 2a). Mastadenovirus showed an uneven genome coverage which could result from contaminating vector sequences. Therefore, we also analyzed the virus detections across sequencing dates (Supplementary Figure 2b) to assess any batch effect indicative of a contanimant; for example in mastadenovirus, we identified an association with sequencing date in early-onset prostate cancer regardless of tumor/normal state. We conclude that our mastadenovirus detections are due to a contamination occurring across projects worldwide as similar patterns could be identified in other projects as well (data not shown).

**Figure 1:**
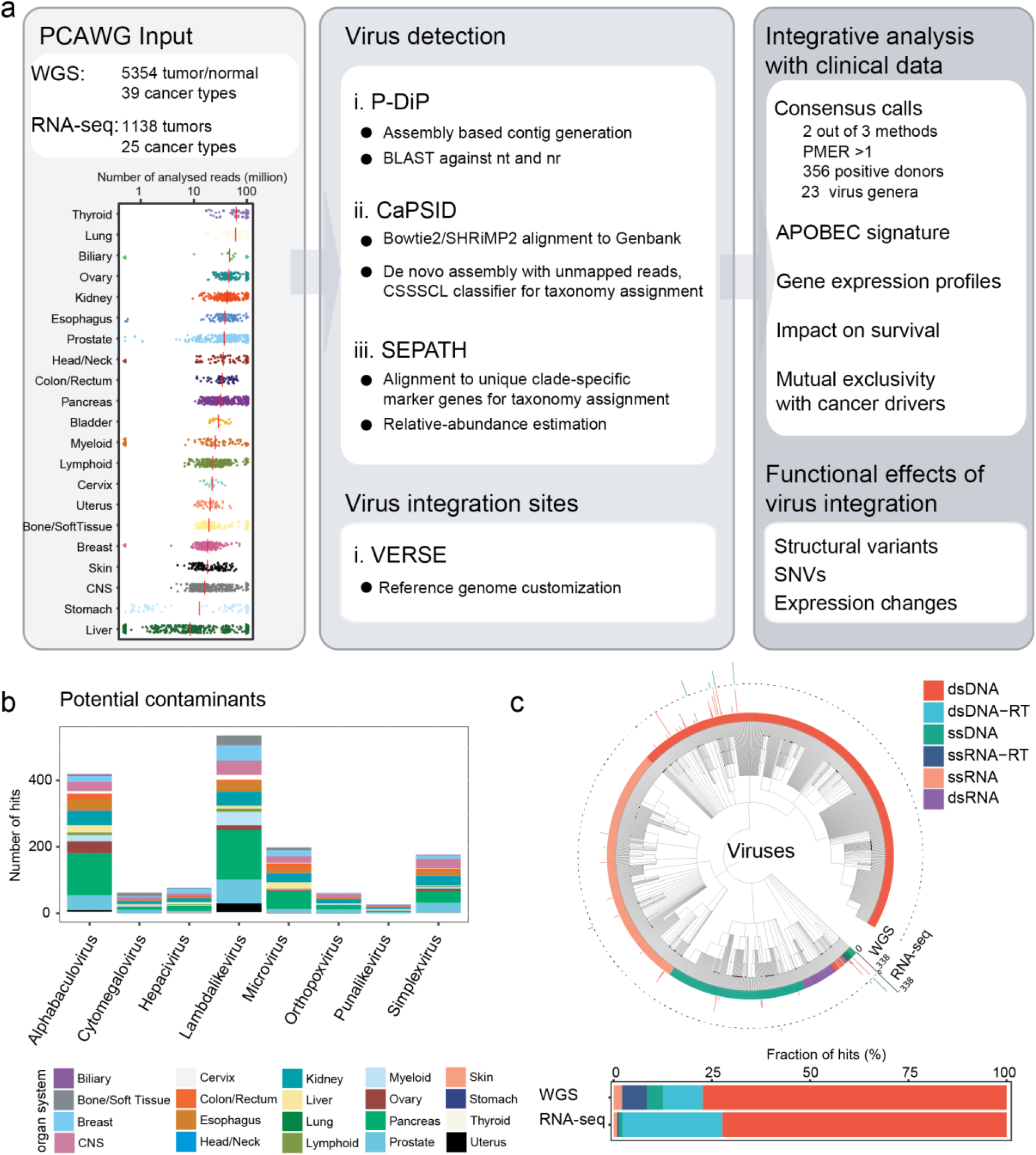
Overview, design and summary statistics. (a) Workflow to identify and characterize viral sequences from the whole-genome and RNA sequencing of tumor and non-malignant samples. Viral hits were characterized in detail using several clinical annotations and resources generated by PCAWG. (b) Identified viral hits in contigs showing higher PMER’s (viral reads **p**er **m**illion **e**xtracted **r**eads) for artificial sequences like vectors than the virus. Displayed are all viruses that occur in at least 20 primary tumor samples in the same contig together with an artificial sequence. (c) Summary of the viral search space used in the analysis grouped by virus genome type. The number of virus positive tumor samples are indicated in the outer rings (PMER log scale for WGS and RNA sequencing data) as detected by any of the pipelines. Taxonomic relations between the viruses are indicated by the phylogenetic tree. dsDNA: double stranded DNA virus, dsDNA-RT: double-stranded DNA reverse transcriptase virus, ssDNA: single-stranded DNA virus, ssRNA-RT: single-stranded RNA reverse transcriptase virus, ssRNA: single-stranded RNA virus, dsRNA: double-stranded RNA virus. Fraction of hits in WGS and RNA sequencing data are depicted as stacked barplot.

We generally observed a strong overlap of the genera identified across pipelines (Supplementary Figure 1d). From the whole genome dataset, we identified 321, 598 and 206 virus-tumor pairs for P-DiP, CaPSID and SEPATH, respectively (Figure 2a, overlap after random permutation of pipeline detections in Supplementary Figure 3a). Notably, there was no difference in the PMER distribution of common hits across the three pipelines indicating that a common detection cut-off is reasonable (Supplementary Figure 2b). The number of hits derived from the RNA-seq dataset differed between the pipelines (positive virus-tumor pairs: 108 for P-DiP, 83 for CaPSID and 41 for SEPATH; Figure 2b). SEPATH, using a k-mer approach, detected the lowest number of virus hits and was the least sensitive. Despite this, the identified viruses matched well with the consensus (DNA 90%, RNA 95%). P-DiP, based on an assembly and BLAST approach detected more hits with 59% of the DNA and 54% of the RNA hits in the consensus set, while CaPSID, being most sensitive, implementing a two-step alignment process complemented by an assembly step, identified 60% (DNA) and 80% (RNA) hits within the consensus set. While the majority of the virus hits from RNA-seq (n=61/68) were overlapping with the WGS data, the reverse is not true, emphasizing the importance of DNA sequencing for generating an unbiased catalogue of tumor-associated viruses. This difference can also be attributed to the viral life cycle as during incubation or latent phases, viral gene expression can be minimal^13^. Contrasting virus positive and virus negative samples within each organ type shows that the organ system as expected has a significant influence (P=<2e-16, ANOVA modeling potential pathogenic reads dependent on organ system and virus positivity, Supplementary Figure 1c) but not virus positivity. This indicates that virus positive tumors are not detected due to a higher number of candidate reads and is in line with the fact that the viral reads in most cases do not substantially contribute to the candidate reads analyzed. 86% of the sequence hits detected from WGS and RNA-seq data were found to be from the virus genome type of double-stranded DNA virus (dsDNA) and dsDNA with reverse transcriptase (Figure 1c). This could be attributed to i) a higher frequency of tumor-associated viruses from these genome types^15^, ii) a larger sequence dataset for WGS in comparison to RNA-seq, iii) a potential limitation of our analysis due to DNA and RNA extraction protocols that are less likely to include ssDNA or RNA viruses or iv) the selection bias of tumor entities included in the PCAWG study (Figure 1c).

**Figure 2:**
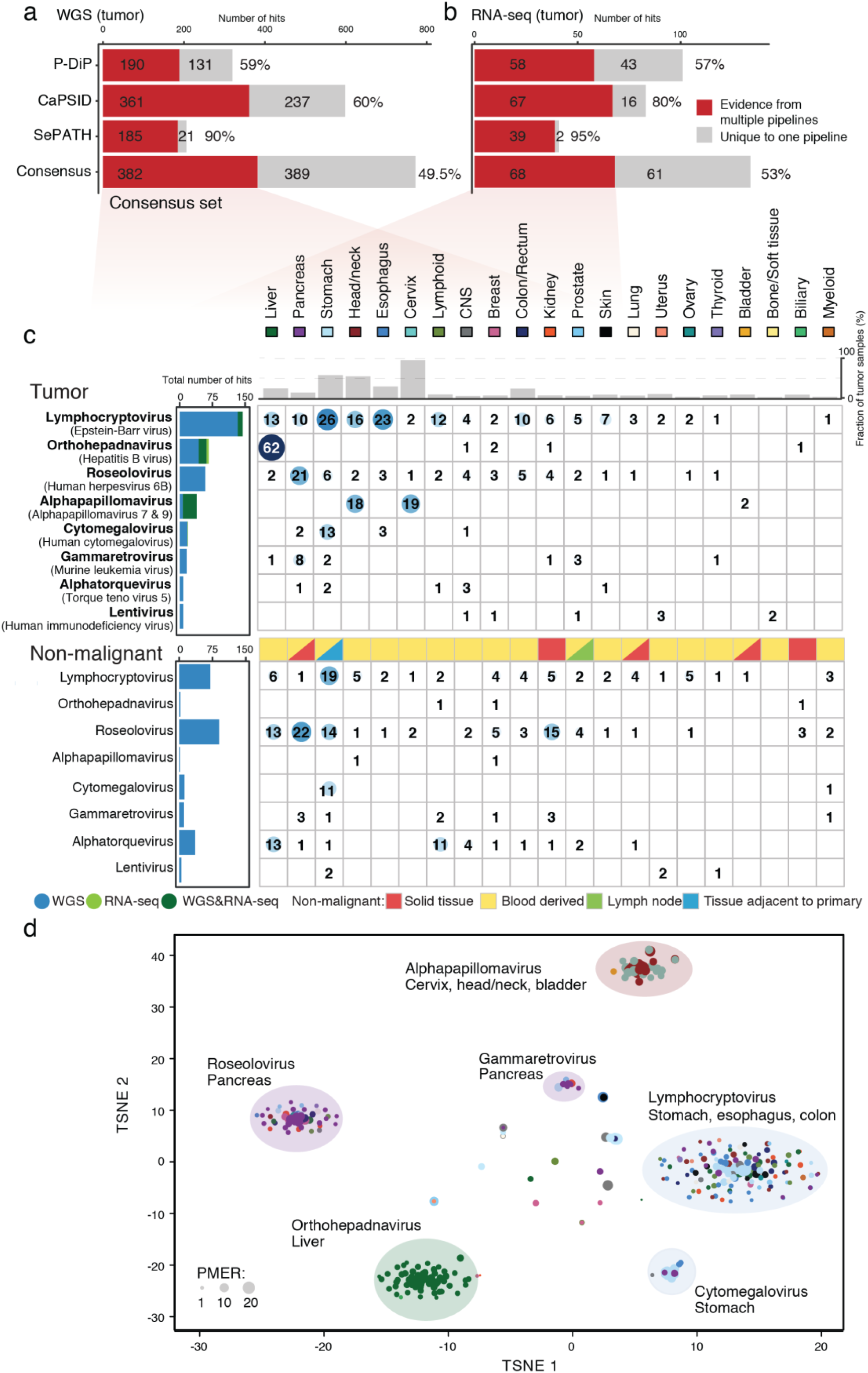
Detected viruses: Consensus for detected viruses in whole genome and transcriptome sequences. Number of genus hits among tumor samples for the three independent pipelines and the consensus set defined by evidence from multiple pipelines. (a) based on whole genome sequencing, (b) and based on transcriptome sequencing. (c) Heatmap showing the total number of viruses detected across various cancer entities. The sequencing data used for detection is indicated among the total number of hits (WGS= blue, RNA-seq=green). The fraction of virus positive samples is shown on top and the type of non-malignant tissue used in the analysis is indicated if more than 15% of the analyzed samples are from a respective tissue type (solid tissue, lymph node, blood or adjacent to primary tumor). (d) t-SNE clustering of the tumor samples based on PMER of their consensus virome profiles, using Pearson correlation as the distance metric. Major clusters are highlighted by indicating the strongest viral genus and the dominant tissue types that are positive in that cluster. Dot size represents the viral reads per million extracted reads (PMER).

### The virome landscape across 39 distinct tumor types

We employed a consensus approach that resulted in a reliable set of 389 distinct virus-tumor pairs from WGS and RNA-seq data (Figure 2, see Materials and Methods). Overall, 23 virus genera were detected across 356 tumor patients (13%). The top five most prevalent viruses (lymphocryptovirus, orthohepadnavirus, roseolovirus, alphapapillomavirus and cytomegalovirus) account for 85% of the consensus virus hits in tumors (n=329 out of 389). Among these five prevalent virus genera, three have been well described in the literature as drivers of tumor initiation and progression^9^: i) lymphocryptovirus (n=145 samples, 5.5%, e.g. Epstein-Barr Virus, EBV) is the most common viral infection across a variety of tumor entities mainly from gastrointestinal tract, and showed a much lower prevalence in the matched non-malignant control samples (n=82, 3%) (Figure 2c); ii) orthohepadnavirus (n=67, 2.5%, e.g. hepatitis B, HBV) are as expected the most frequent among liver cancer with Hepatitis B present in 62 of 330 donors (18.9%); and iii) alphapapillomavirus (findings discussed in detail below). Lymphocryptovirus (n=11), orthohepadnavirus (n=18) and alphapapillomavirus (n=32) were detected both in RNA and DNA sequencing data (Figure 2c, left panel), with Alphapapillomavirus being the most frequent (32 out of 39 consensus hits). This is in line with the constitutive expression of viral oncogenes in cancers associated with these viruses, a parameter supporting a direct role in carcinogenesis^9^. In contrast, our analysis did not find any support at the RNA-seq level for the remaining common genera, such as Roseolovirus. An in-depth analysis of the virus genome equivalents per human tumor genome equivalent considering genome sizes, coverage and tumor purity showed overall low viral genome equivalents even for established tumor viruses (Supplementary figure 3c and Supplementary table sheet “Virus Load”). Evidence for MMTV (PMER = 3.4) was detected in one renal carcinoma sample and in none of the 214 analyzed breast cancer samples. Previous work has suggested that MMTV may play a role in breast cancer but our extensive search of viral sequences could not reveal any MMTV-positive case in breast cancer that would support this claim.

Roseolovirus and Alphatorquevirus show a higher number of hits in the non-malignant control samples, which were mainly derived from blood cells (Figure 2c). For example, we identified 59 patients as Roseolovirus-positive in their tumor and 90 patients positive in the non-malignant control samples. The genus Roseolovirus is composed of human herpesvirus HHV-6A, HHV-6B and HHV-7. Infections occur typically early in life and result in chronic viral latency in several cell types, mainly umbilical cord blood lymphocytes and peripheral blood mononuclear cells^14^. In our systematic study, we detected Roseolovirus mainly in pancreas, stomach and colon/rectum tumors (6%, 8% and 8.3%). Considering the known cell tropism of roseolovirus for B- and T-cells, we asked whether immune infiltration would be higher in roseolovirus-positive tumors. However, we could not identify a stronger contribution of immune cells in virus positive tumor cases as estimated using CIBERSORT^15^ (FDR corrected p-value>0.05 for pancreas; Wilcoxon Rank Sum test for cases with n>3; Supplementary Figure 4a). Therefore, virus positivity cannot be directly linked to immune cell content in the tumor. Still, we cannot rule out a substantial contribution of infected immune cells in pancreas and other tissues. Therefore, in line with current knowledge (reviewed in^16^), we cannot confirm a link between roseolovirus and tumor development. Especially in matched non-malignant kidney samples, we detected higher roseolovirus positivity without an equally strong signal in the corresponding tumor samples. Furthermore, we could not identify actively transcribed viral genes for Roseolovirus and Alphatorquevirus at the transcriptome level. This is in agreement with the latent state of these viruses reported for blood mononuclear cells^14^, and their transmission through blood transfusions (e.g. alphatorquevirus and unclassified anelloviridae^17^). Cytomegalovirus (CMV) was found after identifying and removing contaminations both in stomach tumors (n=13) and the adjacent non-malignant tissue (n=11). CMV is linked to inflammation of the stomach or intestine^18^, as well as infections of the lung and the back of the eye. In line with a recent publication^19^ we could not detect CMV in the analyzed 294 CNS tumors (146 medulloblastomas, 89 pilocytic astrocytoma, 41 glioblastomas, 18 oligodendrogliomas). Therefore, a previously debated role of this virus is not supported. Interestingly, we did not identify a significant enrichment of co-infection of multiple viruses in any tumor type (Supplementary Figure 3d).

#### Hepatitis B virus

Hepatitis B virus was most frequently detected among liver cancers (n=62). Comparing to the histopathological gold standard HBV PCR test^20, 21^ on 228 samples, we found the WGS based consensus detections had the same high specificity (96.1%) and a higher sensitivity (84.0%), indicating that the HBV detections by WGS are reliable (Figure 3a and Supplementary Figure 4b). Furthermore, five out of seven cases positive in WGS and negative for HBV PCR showed positivity for HBAg indicating a high sensitivity of the WGS analysis. In summary, the precision (85.7%) and recall (84%) for the detection of HBV based on ~30x WGS is comparable to targeted PCR. We confirmed a significant exclusivity between HBV infection and CTNNB1, TP53 and ARID1A mutations that was found in a larger liver cancer cohort analyzed by high throughput sequencing (*q*=5.35×10-6, q=0.0023 and q=0.0023, DISCOVER^22^)^23^.

**Figure 3:**
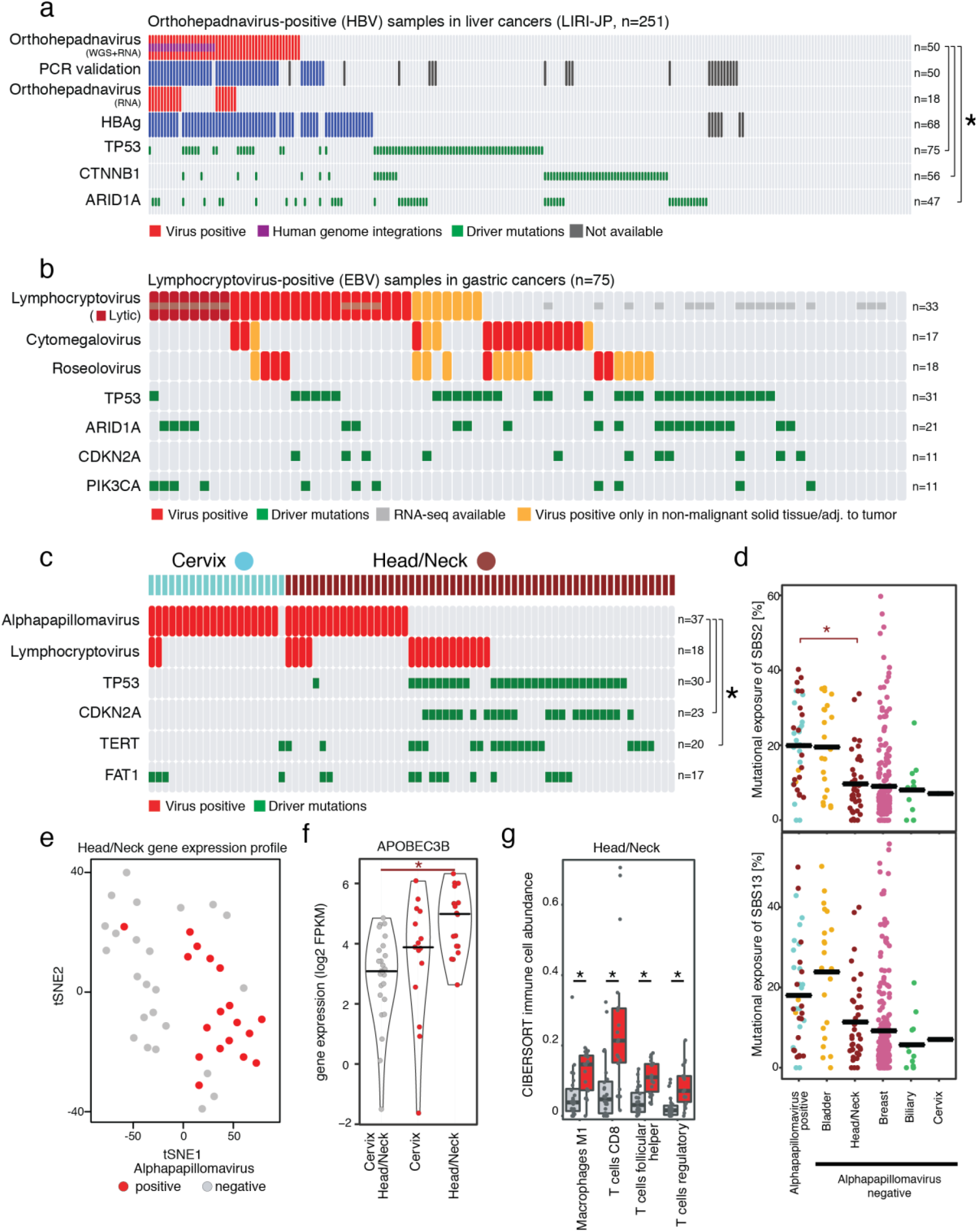
Virus specific findings. (a) Hepatitis B virus detections, validations and driver mutations in liver cancer. Star indicating mutual exclusivity between HBV detections and somatic driver gene mutations. (b) Virus detections in gastric cancer samples, indication of virus phase (lytic/latent) and driver mutations (c) Virus detections and driver mutations in cervix and head and neck cancer. Star indicating mutual exclusivity between alphapapillomavirus detections and somatic driver gene mutations. (d) Alphapapillomavirus detection and exposures of mutational APOBEC signatures SBS2 and SBS13. Star indicates significant difference of mutational signature exposure. (e) Gene expression based tSNE map of head and neck cancer samples show a distinct gene expression profile for virus positive samples. (f) The violin plot of APOBEC3B gene expression for alphapapillomavirus positive and negative samples in cervix and head/neck cancer (Significance of FDR corrected Wilcoxon Rank Sum test is indicated by star) (g) Tumor-infiltrating immune cells as quantified by CIBERSORT linked to alphapapillomavirus infections in head and neck cancer. All four cell types show a significant enrichment of immune cells in virus positive samples (Significance of FDR corrected Wilcoxon Rank Sum test indicated by star).

#### Epstein-Barr virus

Epstein-Barr virus was detected in many different tumor entities and normal samples (Figure 2c). Comparing EBV PMERs in tumors and matched normals we see a stronger contribution in matched normals from matched solid tissue or tissue adjacent to the tumor (Supplementary Figure 4c). For samples showing reads for EBV in WGS and with available RNA sequencing data, the absolute score for immune cells based on CIBERSORT^15^ was not significantly different between virus positive and negative samples (FDR corrected p-value>0.05 for colon/rectum, head/neck, lymphoid, stomach; Wilcoxon Rank Sum test for cases with n>3; Supplementary Figure 4a). In summary, there is no evidence for a detection of EBV due to infiltrating immune cells. This indicates EBV presence in the respective organs. Based on the expression data available for the tumor samples we identified viral transcripts of the latent as well as lytic phase of the viral lifecycle (Figure 3b and Supplementary Figure 3d). Eight of the nine tumors expressing lytic EBV transcripts, are from stomach, confirming its active contribution to stomach cancer^24^.

#### Alphapapillomavirus

Alphapapillomaviruses were mainly detected in head and neck cancer (n=18 out of 57), cervical cancer (n=19 of 20) and in two bladder cancer cases out of 23, in agreement with previous studies^4, 25, 26^. There is also supporting evidence for 32 out of 39 alphapapillomavirus hits in the transcriptome data (Figure 2c). We observed only one HPV subtype per tumor according to the P-DiP results. At the subtype level, HPV16 was found to be the dominant type in cervix (n=11) and head and neck (n=15) tumors, followed by HPV18 only present in cervical cancer (n=6). As reported previously^27^, HPV33 was identified both in head and neck (n=3) and cervix (n=1) tumor samples. Different HPV variants, type 6 and type 45, were detected in bladder cancer.

We further characterized the functional effects of alphapapillomaviruses in tumors by integrating external PCAWG datasets such as driver mutations, mutational signatures, structural variations, gene expression profiles and patient survival. In head and neck cancer, HPV-positive tumors exhibit an almost complete mutual exclusivity with mutations in known drivers like *TP53, CDKN2A* and *TERT* (*q*=1.73×10-5, *q*=1.73×10-5, *q*=0.012; multiple testing corrected for presented mutations and in EBV and HPV, DISCOVER^22^) (Figure 3c), as reported previously^25^, which could be explained by a mutation independent inactivation of TP53 through the human papillomaviruses^28–30^. Analyzing the mutational signatures enriched in these cases, we identified mutational signatures 2 as enriched for alphapapillomavirus positive cases in head and neck cancers (q=0.02; FDR corrected Wilkoxon Rank Sum test; Figure 3d)^31^. In addition, the expression of APOBEC3B is significantly higher in the virus positive head and neck cancers compared to their virus negative counterparts (P<0.001, Wilcoxon Rank Sum Test, Figure 3e)^32^. However, we did not observe the enrichment of APOBEC signatures and expression changes for EBV positive samples neither in cervix nor in other tissues.

Distinct expression profiles between virus positive and negative tumors in head and neck cancer are observed (Figure 3f)^33^. Analyzing the immune cells estimated by CiBERSORT, we could identify a significant increase in macrophages and T-cell signals in alphapapillomavirus positive head and neck cancers (FDR corrected for all viruses and cell types tested, p-values: T cell follicular helper 0.004, T cells CD8 0.012, T cells regulatory 0.012, Macrophages M1 0.018; Wilcoxon Rank Sum test; Figure 3g). Our integrative analysis on HPV reconfirms many of the findings related to HPV infection, illustrating the potential of our systematic approach in identifying and characterizing tumor-associated viruses.

### Transcriptional activation of endogenous retroviruses linked to clinical outcome

Human endogenous retroviruses (HERV) are integrated in the human DNA originating from infection of germline cells by retroviruses over millions of years^34^ and contribute 2.7% of the overall sequence to over 500,000 individual sites in the human genome^35, 36^. The endogenous retroviruses were identified by the three pathogen detection pipelines but filtered by CaPSID and SEPATH. In addition, an alignment-based approach was used to detect HERV sequences embedded in the human reference genome that could be missed by the pipelines focusing only on non-human reads. In this study, we quantified the expression of HERV-like LTR (long terminal repeat) retrotransposons categorized into several clades by Repbase^37^ database as ERVL, ERVL-MaLR, ERV1, ERVK and ERV (Supplementary Table, sheet “HERV expression”). In comparison to the other HERV families, ERV1 shows the strongest expression on average (Figure 4a) and ERVK the highest fraction of active loci (Figure 4b). Analyzing the expression of HERVs based on the available RNA sequencing data, we could identify a strong expression for ERV1 in chronic lymphocytic leukemia compared to all other tumor tissues and adjacent normal tissues (Figure 4c). However, we could not identify a link between transcriptionally active stemness markers (OCT3/4, SOX2, KLF4) and increased HERV expression as opposed to Ohnuki et al. ^38^(Spearman Rank correlation < 0.35, Supplementary Figure 5). New data suggest that expression of HERVs is associated with prognosis in clear cell renal cell carcinoma (ccRCC)^39^. Analyzing the HERV expression in relation to patient survival, we identified a high ERV1 expression in kidney cancer linked to worse survival outcome (*P* = 0.0081; Log-rank test; Figure 4d, for other HERVs and tumor types see Supplementary Figure 6).

**Figure 4:**
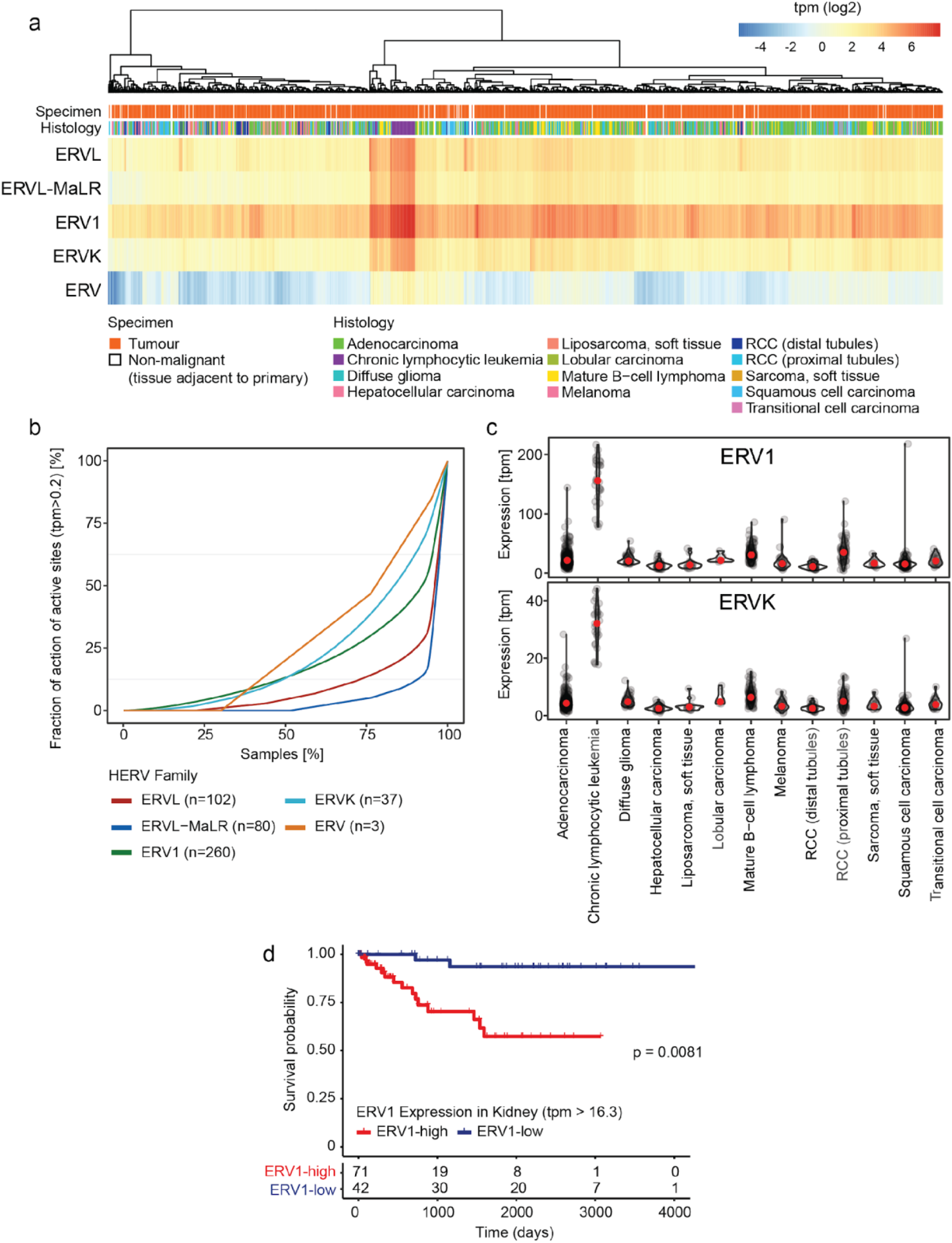
Endogenous retroviruses. (a) Heatmap showing the HERV expression across all tumor samples. HERV TPMs were grouped by family and summed up. Hierarchical clustering was performed by family based on Manhattan distance with complete linkage after log2 transformation of HERVs TPM expression values. (b) Fraction of active loci in the genome with a TPM >0.2 plotted against the fraction of samples. (c) TPM based expression of the highly expressed HERVs ERV1 and ERVK across tumor types. (d) Survival difference between kidney cancer samples expressing high and low levels of ERV1.

### Genomic integration of viral sequences

Viral integration into the host genome has been shown to be a causal mechanism that can lead to cancer development^40^. This process is well-established for human papilloma viruses (HPVs) in cervical, head-and-neck and several other carcinomas, and for hepatitis B virus (HBV) in liver cancer^41, 42^. We searched the PCAWG genome and transcriptome cohorts for integration of those viruses that were detected by the CaPSID platform using the “Virus intEgration sites through iterative Reference SEquence customization” (VERSE) algorithm^43^. This algorithm utilizes chimeric paired-end as well as soft-clipped sequence reads to determine integration with single base-pair resolution. Detailed assessment of this algorithm (e.g. distinction from background noise) is presented in the methods section (see Materials and Methods).

Low confidence integration events were detected for the two viruses HHV4 (in gastric cancer and malignant lymphoma) and HPV6b (head and neck and bladder carcinoma), while integration events with high confidence were demonstrated for HBV (liver cancer), Adeno-associated virus-2 (AAV2) (liver), HPV16 and HPV18 (both in cervical and head and neck carcinoma). Most of these integration events were found to be distributed across chromosomes and a significant number of viral integrations occur in the intronic (40%) regions while only 3.4% were detected in gene coding regions (Supplementary Figure 7a-d).

HBV was found to be integrated in 36 liver cancer specimens out of 61 patients identified as HBV-positive. Notably, genomic clusters of viral integrations (see Materials and Methods) were identified in *TERT* (ngc = 6, where ngc indicates the number of integration sites within a genomic cluster), *KMT2B* (ngc = 4), recently identified to be a likely cancer driver gene^44, 45^ and *RGS12* (ngc = 3) (Supplementary Figure 7e). Furthermore, two or more integration events in individual samples were observed in the gene (or gene promoter) regions of *CCCNE1, CDK15, FSIP2, HEATR6, LINC01158, MARS2* and *SLC1A7* (Figure 5). Additional events with two integration sites were also detected within a 50 kb distance away from *CLMP, CNTNAP2* and *LINC00359* genes. Integration events at *TERT* were found to recur in five different liver cancer samples. One sample had a genomic cluster of three viral integration events within *TERT* and four samples contained a single integration event in the *TERT* promoter (3) or 5’ UTR regions (Supplementary Table Sheet “Integration”). When comparing gene expression in samples with virus integration to those without, only TERT was over-expressed (fold change ≥ 2.0) in two liver cancer samples (Figure 5e). Additional genes with increased expression impacted by integration events include *TEKT3, CCNA2, CDK15* and *THRB* (Figure 5a).

**Figure 5:**
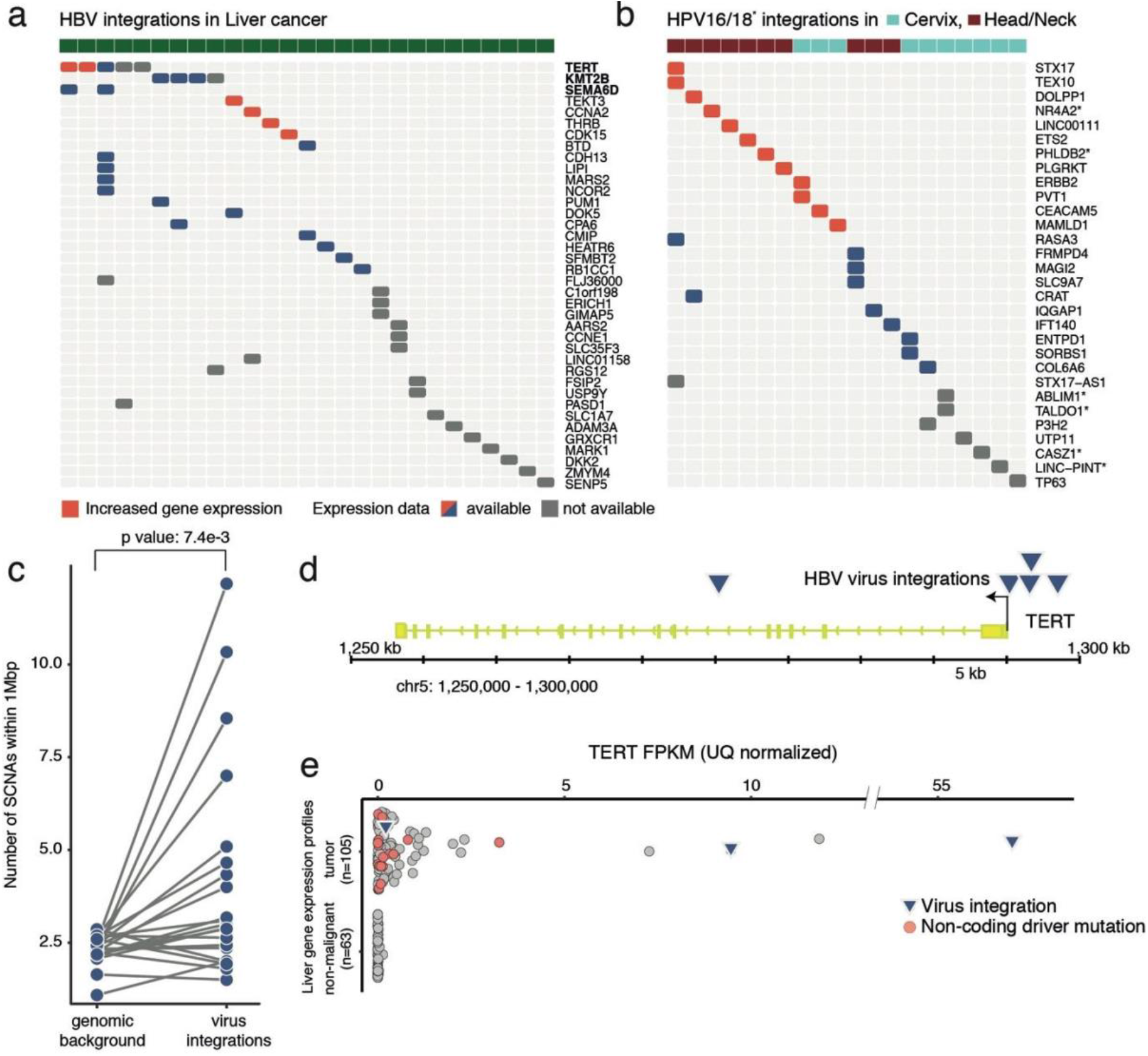
Impact of virus integration. (a) Integration sites detected in gene regions (including promoter, exon, intron and fiveprimeUTR regions) are labeled in red for increased gene expression and blue for expression measured. Rows of each heatmap designate nearest genes to the integration sites and columns represent individual ICGC donor and project ids. Intragenic HBV integration sites detected in liver cancers (ICGC project codes: LIRI, LIHC and LINC). For TERT and SEMA6D intergenic integrations are shown as well. (b) Integration sites detected for HPV-16 and 18 in head/neck (samples color coded magenta) and cervical (samples color coded blue) cancers (ICGC project codes: HNSC and CESC) gene labels with star indicated HPV18 as opposed to HPV16 viral integrations. (c) A local increase in the number of SCNAs was shown in the vicinity of HBV viral integrations (n=21). (d) Genomic visualization of the HBV virus integration sites relative to the TERT gene in five liver tumor patients. (e) The increased gene expression (FPKM) of TERT gene in two liver tumors with HBV viral integrations in comparison to the TERT expression in tumor and non-malignant adjacent tissue. Tumor samples with a non-coding driver mutation were labeled in orange.

Novel genes, which are impacted by integration events and associated with cancer include: *CDK15* that was found to be over-expressed in our study and reported to mediate resistance to the tumor necrosis factor-related apoptosis-inducing ligand (TRAIL)^46^; *SEMA6D* identified as a potential oncogene candidate in human osteosarcoma^47^; and *CDH13* that is commonly downregulated through promoter methylation in various cancers^48^. In addition, a novel integration site located in the promoter region of *ERICH1* was detected (Supplementary Table, Sheet “Integration”).

There was a significant association between HBV viral integrations and SCNAs (Figure 5c). For samples with HBV integration events, the number of SCNAs was higher on average in the vicinity of viral integration sites (within 1 Mbp) when compared to samples without HBV integration (4.2 vs 2.3, *P* = 7.0×10-3; two-sided paired *t*-test). No evidence for an SCNA association was seen for other integrated viruses like HPV16/18 (Supplementary Figure 8a-b).

HPV18 integration events were detected in seven tumors in total, with the most notable clusters of integration events in cervical cancer samples affecting *TALDO1* (ngc = 4) (Supplementary Figure 7g). As shown in Figure 5b, single viral integration events were detected in the genes *CASZ1, LINC-PINT, NR4A2, ABLIM1* and *PHLDB2*.

In 20 samples, HPV16 integration events were detected. Genomic clusters of viral integration sites were identified in cervical and head and neck cancer samples affecting the genes *PVT1* (ngc=9), *PLGRKT* (ngc=4), *ETS2* (ngc=3), *LINC00111* (ngc=3) and *TEXT10* (ngc=3) (Supplementary Figure 7f). Additional integration events with at least two sites were detected in *CRAT, ERBB2, FRMPD4, MAGI2, MAMLD1, SLC9A7, STX17* and *TP63*. None of these multiple integration events were observed to recur across multiple patients (Figure 5b). Integration events were also observed in two different lncRNAs, the plasmacytoma variant translocation 1 gene (*PVT1*), which is recognized as an oncogenic lncRNA observed in multiple cancers including cervical carcinoma^49, 50^, and *LINC00111*, the function of which is still to be determined. Expression of both genes is strongly increased in the cases with HPV16 integration (Supplementary Figure 8f). Individual HPV16 integration sites were also found in a number of other genes including known drivers of tumor pathogenesis (*TP63, P3H2, ETS2, CD274* (PD-L1), *ERBB2, IQGAP1*) (see Supplementary Tables, sheet “Integration”) and genes that were previously not known to be strong candidates for playing a role in tumorigenesis (*CEACAM5, CRAT, ENTPD1-AS1, FRMPD4, MAGI2, MAMLD1, UTP11, COL6A6, RASA3, SORBS1, STX17-AS1, IFT140 and DOLPP1*).

Using the merged single nucleotide variant (SNV) calls from the three mutation calling pipelines (DKFZ, Broad and Sanger)^10^, and by comparing samples with viral integration events to those without, we have found a significant increase in the number of mutations occurring within +/- 10,000 bp of high-confidence viral integration sites (average number of mutations per sample = 0.41 (HPV16 +) vs 0.14 (HPV16 -), *P* = 0.02; paired t-test one sided - alternative greater, Supplementary Figure 8 c, d). Interestingly the integration sites are, compared to a random genome background, enriched in close proximity (<1000bp) to common fragile sites (*P* = 0.0018, two sided Kolmogorov–Smirnov test). These results suggest that HPV16 integration reflects either characteristics of chromatin features that favor viral integration, such as fragile sites or regions with limited access to DNA repair complexes, or the influence of integrated HPV16 on the host genome, both in close vicinity and a long distance away from the integration site. Such a correlation was not seen for the integration sites of other viruses (see Supplementary Figure 8e).

Finally, a single AAV2 integration event located in the intronic region of the cancer driver gene KMT2B^51^ was detected in one liver cancer sample.

### Identification of novel viral species or strains

The CaPSID pipeline, combining both the reference based and de novo assembly approach, was used to search for potentially novel virus genera or species. De novo analysis has generated 56 different contigs that have been classified into taxonomic groups at the genus level by the CSSSCL algorithm^52^. After filtering de novo contigs for their homology to known reference sequences, we have identified 29 contigs in 28 different tumor samples showing low sequence similarity (in average 63%) to any nucleotide sequence contained in the Blast database (see Materials and Methods). In this respect, our analysis has shown that WGS and RNA-seq can be used to identify novel isolates potentially from new viral species. However, the total number of novel isolates were quite low in comparison to viral hits to well-defined genera (Figure 2c). These *de novo* contigs were not enriched for a specific tumor entity but rather distributed across cancer types including bladder, head/neck and cervical cancers and more (Supplementary Figure 9).

## Conclusions

Searching large pan-cancer genome and transcriptome data sets allowed the identification of an unexpectedly high percentage of virus associated cases (16%). In particular, analysis of tumor genomes, which were sequenced on average to a depth of at least 30 fold coverage, revealed considerably more virus positive cases than investigations of transcriptome data alone, which is the search space looked at in most previous virome studies. This is probably mainly due to viruses with no or only weak transcriptional activity in the given tumor tissue. Co-infections, generally believed to indicate a weak immune system, were very rare (Supplementary Figure 3d). This could, however, also be the result of selection processes during tumorigenesis.

While universal criteria for a causality of viral pathogens are prone to errors, it is worthwhile to look at individual features that might support a potentially pathomechanistic contribution of a given pathogen. These include aspects that affect the expression of host factors, e.g. upon viral integration, or the mutual exclusivity of the presence of viral genomes and other host factors, which are already known to play a role in the etiology of a given tumor type. Such aspects need to be carefully considered when discussing of what strengthens a potentially pathogenic role of virus.

Not surprisingly, known tumor associated viruses, such as EBV, HBV and HPV16/18, were among the most frequently detected targets. Interestingly, viral detection based on whole genome sequencing showed similar performance with respect to precision and recall as a targeted PCR for HBV indicating the sensitivity of this approach to detect viruses. This is in particular true for the common integration verified for HBV and HPV 16/18 in our study. In addition, the common theme of potential pathomechanistic effects by the genomic integration of viruses, nurtured also by the observations of multiple nearby integration sites in a given tumor genome that we also report in the present study, has gained further momentum. Analyzing the effect of viral integrations on gene expression, we identified several links to genes nearby the integration site. In this regard, the frequently observed integration of HBV at the *TERT* promoter accompanied with the transcriptional upregulation of *TERT*, constitutes an intriguing example, since an increased activity of TERT is a well-understood driver of cancerogenesis^53^. Furthermore, we also linked viral integrations to increased mutations (SNVs and SCNAs) nearby the integration site.

The known causal role of HPV16/18 in several tumor entities, that triggered one of the largest measures in cancer prevention, has been the reason for extensive elucidation of the pathogenetic processes involved. Nevertheless, comprehensive analyses of WGS and RNA-seq data sets revealed additional novel findings. While we confirmed the exclusivity of HPV infection and *TP53, CDKN2A* and *TERT* mutations in head and neck tumors, we could also link virus presence to an increase in mutations attributed to the mutational signature 2^54^. These are explained by the activity of APOBEC, which – among other effects – changes viral genome sequences as a mechanism of cellular defense against viruses^55, 56^. This activation could play an important role in introducing further host genome alterations and, thus, constitute an important mechanism driving tumorigenesis^32, 56^. In liver cancer mutations in *CTNNB1, TP53* and *ARID1A*, major primary oncogenes in this cancer type and HBV infections were confirmed to occur significantly exclusive^23^. Furthermore, the virus positive head and neck cancer samples had a significantly higher abundance of T-cell and M1 macrophage expression signals, which matches with the recently described subtypes of HNSCC that differ – among others – in virus infection and inflammation features.

## Acknowledgments

We thank the IT Core Facility at the DKFZ for technical assistance, as well as to Michael Hain and Rolf Kabbe for computational support. We thank Sabrina Gerhardt for technical assistance in validation experiments. We thank the ICGC/TCGA Pan-Cancer Analysis of Whole Genomes Network.

## Funding

V.F. and I.B. received support for their work from the Ontario Institute for Cancer Research (OICR) through funding provided by the government of Ontario. A.G. received support for his work from the Leibniz Association (Grant Number: SAW-2015-IPB-2) and the German Center for Infection Research (Grant Number: TTU 01.801). P.L. and A.G. received support for this work from the German Federal Ministry of Education and Research (BMBF BioTop Grant Number 01EK1502C, ICGC-DE-Mining Grant Number 01KU1505A-G). D.S.B. received support from Cancer Research UK C5047/A14835/A22530/A17528, the Dallaglio Foundation, Bob Champion Cancer Trust, The Masonic Charitable Foundation successor to The Grand Charity, The King Family and the Stephen Hargrave Trust. H.M. was supported by a Swiss National Science Foundation grant (No. S-87701-03-01).

## Competing interests

The authors have declared that they have no competing interests.

## Notes

#### Summary of Updates

We during the revision process performed detailed analyses and now feel there is insufficient evidence to score mastadenovirus as a tumor relevant event. We included additional analyses on virus genome coverage and sequencing batch effects that strengthen our consensus findings. While the comprehensive analyses could confirm known tumor viruses we make a major contribution about controversial associations of viruses with various tumor entities, as intensely debated in the literature. Despite the comprehensive data set, we could not detect a link of cytomegalovirus with brain cancers and MMLTV with breast cancer.

